# GROMODEX: Optimisation of GROMACS Performance through a Design of Experiment Approach

**DOI:** 10.1101/2025.10.08.681202

**Authors:** Marco Savioli, Paolo Calligari, Ugo Locatelli, Gianfranco Bocchinfuso

## Abstract

We introduce GROMODEX, a novel tool designed to optimise GROMACS molecular dynamics (MD) simulations using a structured Design of Experiments (DoE) approach. GROMACS, though efficient, requires extensive tuning of parameters to perform optimally on different hardware and molecular systems. Manual tuning is tedious and prone to errors, especially for high-performance computing (HPC) environments.

GROMODEX automates this process, systematically exploring performance parameters to find the best configurations for each system. By using statistical techniques, it evaluates multiple parameters simultaneously, optimising for CPU and GPU architectures, and reducing simulation times while ensuring efficient resource use.

We evaluated GROMODEX on four computational platforms spanning a wide range of cost and hardware complexity using four molecular systems from the NVIDIA GROMACS benchmark suite. Quite surprisingly, our results show that the performance does not always correlate with the hardware complexity and cost in a system-dependent manner. Furthermore, a fine system-dependent tuning assures substantial performance gains and time reductions, highlighting GROMODEX’s ability to improve the performance of MD simulations significantly.

## 1. INTRODUCTION

In the realm of MD simulations, GROMACS^1,2^ has firmly established itself as one of the most powerful and widely used tools for simulating molecular systems at the atomic level. Initially developed for biomolecular simulations of proteins, lipids^3^, and nucleic acids, it has expanded its capabilities to simulate diverse molecular systems, including polymers^4,5^ and inorganic materials^6^. However, this versatility comes with the challenge of performance optimisation, especially in high-performance computing (HPC) environments.

One of GROMACS’ key strengths lies in its ability to scale efficiently across CPUs and modern GPU-accelerated systems, offering substantial speedups through parallelisation. Nonetheless, achieving optimal performance is complex, as it depends not only on hardware configurations but also on the system complexity and simulation-specific parameters, such as domain decomposition^7^, Particle Mesh Ewald (PME)^8,9^ settings, as well as the balance between bonded and non-bonded interactions. A poor configuration of these parameters can introduce inefficiencies, and even small parameter adjustments can lead to significant performance changes, making systematic tuning critical.

The diversity of HPC architectures adds further complexity. Hybrid CPU-GPU systems require careful workload balancing to avoid bottlenecks, while factors like memory bandwidth, I/O latency, and interconnects can influence simulation throughput. Manual tuning in such contexts is not only time-consuming but also error-prone, often leading to wasted computational resources. This has sparked interest in automated tools^10^ that streamline optimisation and ensure efficient resource use.

Performance optimisation in MD simulations takes shape of a multidimensional challenge. Historically, optimisation has often been approached using the one–factor–at–a–time (OFAT)^11^ method, which alters a single variable while keeping all others constant. While offering some insights, OFAT has serious limitations in complex multifactorial systems like GROMACS where numerous parameters interact.

By contrast, the DoE methodology^12^ provides a robust and efficient framework for performance optimisation. Unlike OFAT, which assumes independence between parameters, DoE explores multiple parameters simultaneously, revealing complex, nonlinear interactions that often occur in systems like MD simulations. This approach constructs an experimental matrix with predefined levels, reducing the number of trials needed while offering a comprehensive understanding of the parameter space. By analysing the results, DoE not only identifies optimal settings but also uncovers how interdependent variables influence performance, thus enabling more effective configurations.

This approach has proven highly successful across various fields, where multifactorial systems are the norm. In the pharmaceutical industry, it is frequently used to optimise process conditions^13,14^ and analytical method development and validation^15^. It is also extensively used in food industry^14,16^, material and design industry^17^ and environmental chemistry^18^. In all these domains, DoE consistently outperforms OFAT, particularly in cases where factor interactions are critical to optimisation.

The advantages of DoE are particularly compelling in the context of HPC and MD simulations, where the vast number of variables and possible configurations is overwhelming.

Despite its success in other domains, DoE remains underutilised in the optimisation of MD simulations^19^. Currently, most GROMACS users rely on manual tuning or *ad hoc* methods, which are inherently limited by time constraints and the complexity of the parameter space. There is a clear need for a dedicated tool that systematically applies DoE to optimise GROMACS performance across different HPC platforms: a gap that GROMODEX seeks to address.

GROMODEX approaches GROMACS performance optimisation as a well-structured experiment, akin to a scientific study conducted not *in vitro*, but *in silico*. Just as chemists design-controlled experiments to isolate specific effects, GROMODEX systematically explores the performance landscape of a MD simulation. By applying DoE principles, it assesses a wide range of configurations, identifying not only the best settings for a given simulation on a specific machine but also how those settings may generalise across other systems. This makes GROMODEX a valuable tool for researchers working with diverse molecular systems and HPC platforms. This structured approach is a significant departure from traditional trial-and-error methods. By bringing the rigor of experimental design to MD simulation optimisation, GROMODEX transforms performance tuning into a scientifically grounded, reproducible and data-driven process. Moreover, GROMODEX significantly reduces the time and computational resources needed for performance optimisation. In large-scale projects, even small improvements in efficiency can translate into substantial time and cost savings. By automating the optimisation process, GROMODEX allows researchers to focus on scientific inquiries rather than technical fine-tuning.

Lastly, as demand for HPC grows in fields such as computational chemistry, the environmental impact of these platforms is becoming a key concern^20,21^. HPC infrastructures have substantial carbon footprints due to their high energy demands^22–24^.Given the global need to reduce carbon emissions, there is increasing focus on making HPC not only faster but also more sustainable.^25,26^ By enabling GROMACS to use computational resources more efficiently, GROMODEX reduces the overall energy consumed per simulation, helping to lower the environmental impact of MD research. Optimising performance has a direct correlation to reducing energy usage. By maximising throughput and minimising inefficiencies, GROMODEX enhances both speed and energy efficiency. This improvement in computational efficiency has far-reaching implications, enabling more research to be conducted with the same energy input.

By moving away from the traditional OFAT approach and adopting a more holistic DoE framework, GROMODEX offers a deeper understanding of complex parameter interactions and provides a more efficient path to optimisation. With the widespread use of GROMODEX, we aim to establish new best practices in the field, making it a valuable resource for HPC users. As MD simulations continue to grow in scale and complexity, GROMODEX will play an increasingly crucial role in maximising the potential of HPC resources while promoting ecological sustainability.

## 2. METHODS

### 2.1. GROMACS Setup and computational resources

To conduct a comprehensive evaluation of the effectiveness of GROMODEX across a diverse array of hardware infrastructures, MD simulations were carried out on four distinct computational platforms. In all the cases, the performances were registered in the absence of other running processes. These platforms were carefully selected to encompass a broad spectrum of computational capabilities, ensuring a thorough assessment of GROMODEX’s adaptability and performance under varying conditions. The HPC systems involved for this study include:

- **Majorana**: A commodity workstation accessible within our facilities, serving as a baseline for moderate-scale simulations.
- **Franklin**: A high-performance workstation featuring an updated architecture, representing a more advanced yet still locally accessible computational resource.
- **Ada**: A dedicated HPC node within our departmental facilities, providing enhanced processing power and optimised parallel execution.
- **Leonardo**: A state-of-the-art HPC system hosted at CINECA, equipped with cutting-edge computational resources and designed to handle large-scale scientific workloads. It ranks fourth on the TOP500 list^27^ Reported data for Leonardo refer to performance on a single node.

**Table 1** summarises the specifications of the computational platforms used in the present study. Their diversity enables a rigorous evaluation of GROMODEX’s performance, particularly on GPU-accelerated architectures. Majorana, Ada, and Franklin, with multiple GPUs, support the analysis of moderate-scale parallel processing, while Leonardo, with four GPUs, allows us to examine large-scale computational efficiency and performance scaling. This selection ensures a comprehensive assessment of hardware-specific optimisations and parallelisation strategies, making our findings broadly applicable across different computational setups.

**Table 1.**
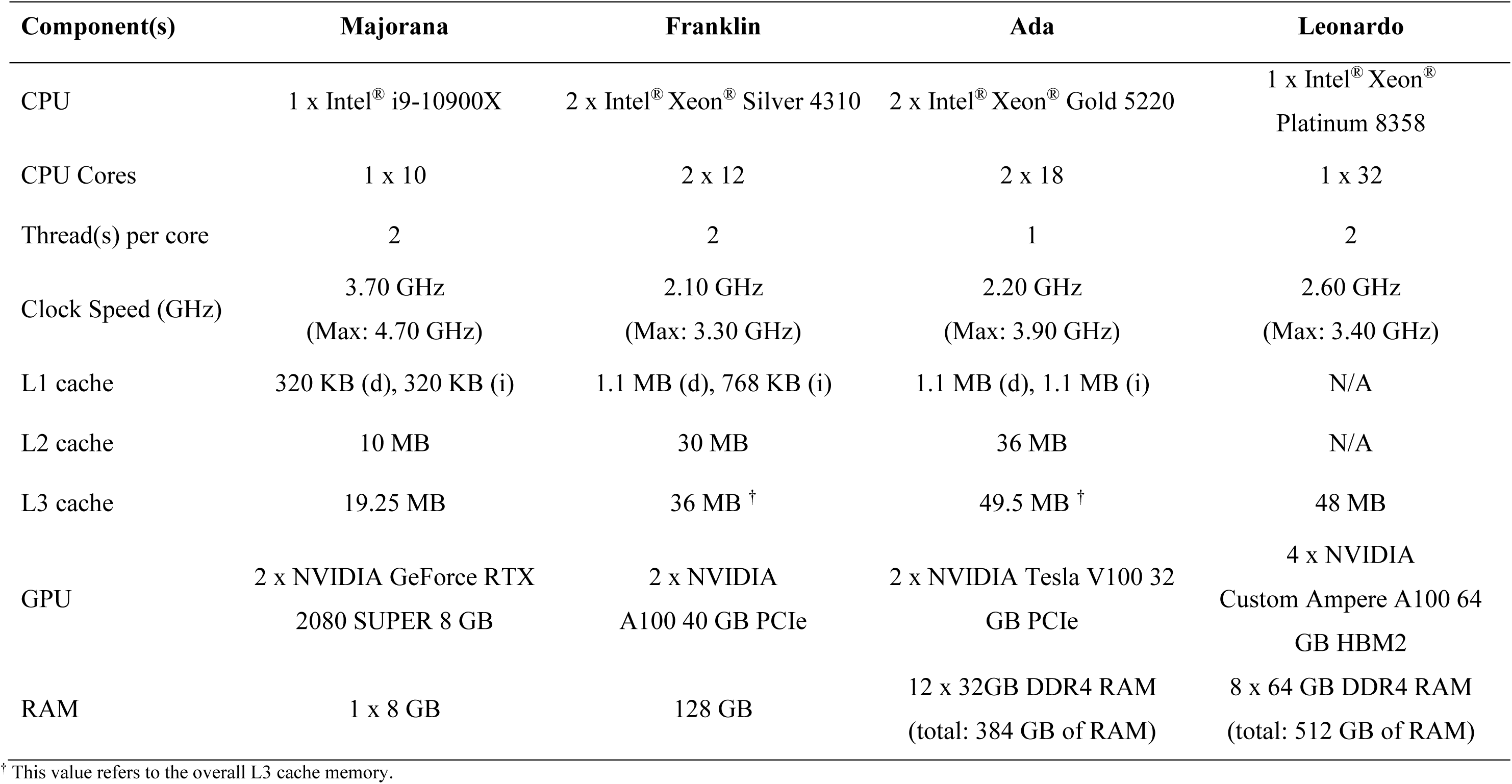
HPC configuration evaluated in the present study.

### 2.2. Molecular Systems and Simulation Parameters

To ensure the robustness and validity of our performance assessment, we employed GROMACS version 2022.6, a highly optimised and widely used MD package. The selection of this software was motivated by its extensive adoption in computational biophysics, its sophisticated parallelisation capabilities, and its compatibility with a range of hardware architectures.

For benchmarking purposes, we selected four molecular systems from the NVIDIA GROMACS benchmark suite^28^, each representing a different level of complexity and system size. These are:

- **Villin**: A small and relatively simple protein, often used as a standard benchmark for MD simulations.
- **RNAse Cubic**: A mid-sized molecular system that introduces moderate computational demands.
- **Alcohol Dehydrogenase (ADH Dodec)**: A larger biomolecular complex, offering a more computationally intensive challenge.
- **Satellite Tobacco Mosaic Virus (STMV)**: A highly complex and large-scale molecular system, representing the upper end of computational feasibility within modern HPC environments.

By selecting systems with varying sizes and levels of complexity, we ensured that our results would be broadly applicable across the typical range of MD simulations encountered in biophysical research. **Table 2** presents an overview of the molecular parameters associated with each of these systems, paralleling those indicated in the NVIDIA benchmark webpage^28^.

**Table 2.**
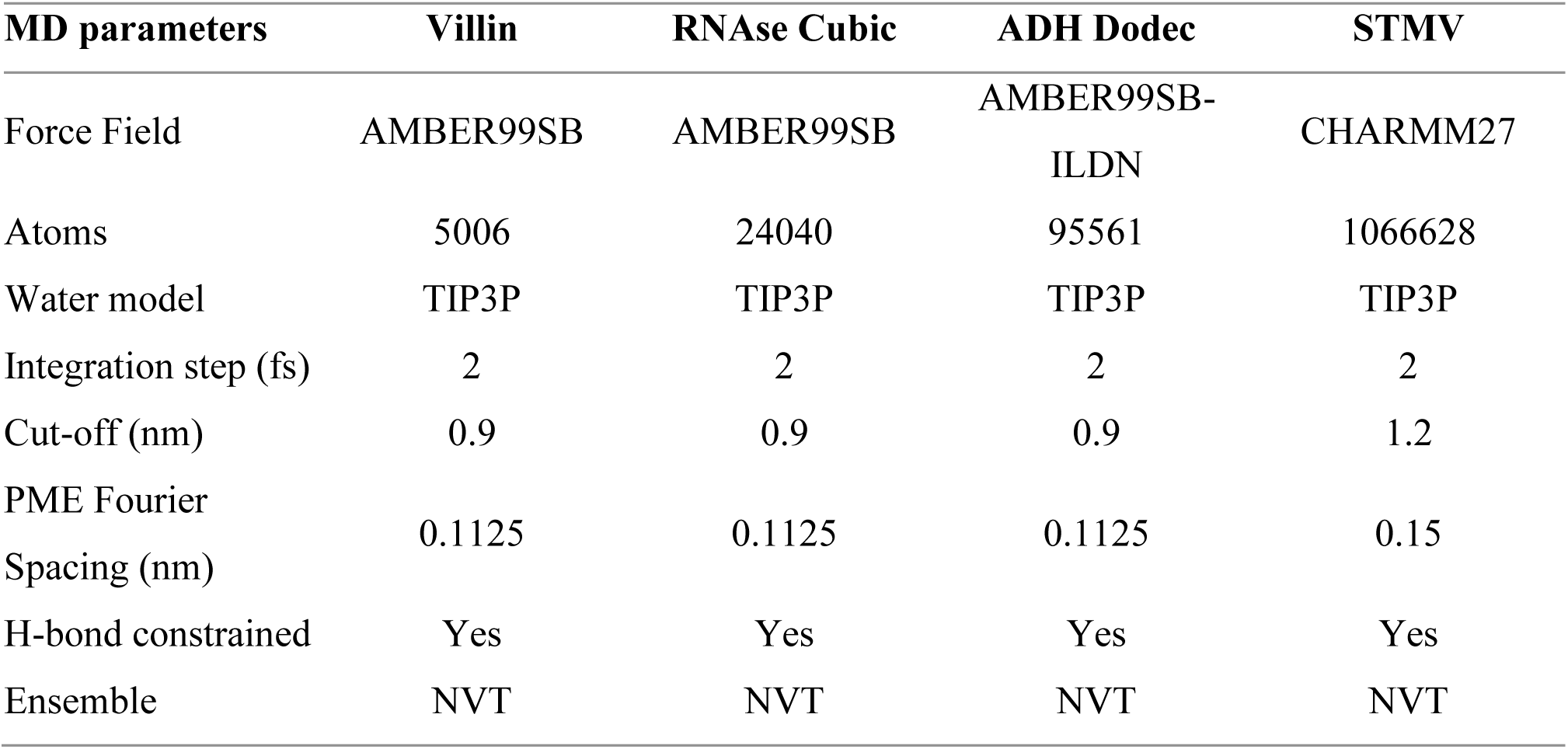
Overview of the molecular system used as performance benchmarks.

### 2.3. DoE Approach

The DoE framework provides a rigorous statistical methodology for systematically analysing the effects of multiple variables on a target response. In computational scenarios such as the performance optimisation of MD engines like GROMACS, DoE enables a structured and efficient exploration of the parameter space, elucidating the individual and joint influence of parameters on key performance indicators.

In this study, we employed a full factorial design, one of the most exhaustive and informative experimental strategies. This approach evaluates all possible combinations of factor levels, offering a complete and unbiased mapping of the experimental domain. When the number of factors and associated levels is tractable, the full factorial design is particularly advantageous as it allows for precise estimation of both main and interaction effects, without the risk of confounding.

The model structure is contingent mainly upon the nature of the response variable, whether categorical or continuous.

Incorporating categorical parameters within a full factorial scheme offers multiple benefits in performance tuning contexts:

- It ensures comprehensive coverage of the configuration space, avoiding bias introduced by heuristic or sequential designs.
- It enhances the detection of synergistic or antagonistic interactions between parameters, often overlooked by OFAT or fractional factorial designs.
- It establishes a solid foundation for subsequent predictive modelling, such as linear regression or surrogate models based on machine learning techniques.

### 2.4. Experimental Factors and Levels

The following factors were chosen as variables in the DoE (Design of Experiments) setup, as they are known to significantly affect the performance of GROMACS simulations across different hardware configurations:

- **ntmpi**: Number of MPI threads per rank (5 levels for two GPUs, 15 levels for four GPUs) – This factor controls the parallelization strategy for message passing, which is essential for scaling simulations across multiple processors. The levels represent varying degrees of parallelism, allowing for a detailed analysis of the impact on performance as the number of threads per rank increases.
- **pin**: CPU core pinning strategy (2 levels) – CPU core pinning refers to the practice of binding specific threads to specific CPU cores to optimise memory locality and reduce inter-core communication overhead. The two levels represent distinct pinning strategies, one where threads are allowed to float between cores, and another where they are pinned to fixed cores, ensuring consistent memory access patterns.
- **gputasks**: Distribution of tasks across GPUs (15 levels for two GPUs, 31 levels for four GPUs) – This factor governs how computational tasks are assigned to GPUs in a multi-GPU setup. The levels vary from distributing tasks evenly across GPUs to more complex configurations where certain GPUs are dedicated to specific types of calculations. This distribution is crucial for optimising GPU utilization and minimizing idle times.
- **bonded**: Bonded interaction calculation method (2 levels) – This factor influences the method used for calculating bonded interactions in MD simulations. The two levels represent different hardware (CPU or GPU) for executing the bonded interactions.
- **update**: Update method for particle positions (2 levels) – The method by which particle positions are updated during the simulation affects the accuracy and efficiency of the calculations. As in the case of the aforementioned parameter, the two levels identify the hardware to set where to execute update and constraints.

Although some of these parameters (such as *ntmpi* and *gputasks*) assume numeric values, all factor levels in this experiment are treated as categorical variables, meaning that the analysis does not rely on any assumed ordering or spacing between the levels.

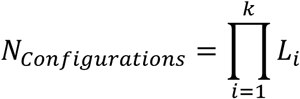

Where:

- *k* = 5 is the number of factors of the experiment,
- *L*_*i*_ is the number of levels of the *i*-th factor.

For HPC systems equipped with 2 GPUs—such as Majorana, Ada, and Franklin—the levels for each factor were defined as:

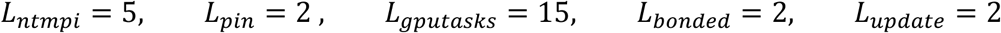

Yielding:

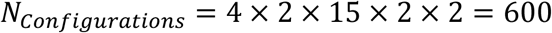

For 4-GPUs HPC systems *gputasks* factor was extended to reflect the increased parallelism complexity and task distribution possibilities. The levels for each factor were defined as:

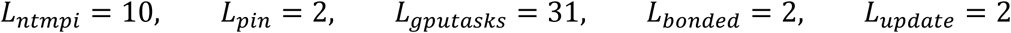

Thus, the total number of unique simulation configurations is:

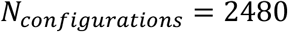

Therefore, 600 (2480 in the case of Leonardo) distinct simulation configurations were systematically generated and executed across different computational platforms, enabling a comprehensive analysis of how performance scales with respect to both software-level parameters and hardware-specific characteristics.

By its nature, a full factorial DoE systematically explores the performance space of the selected parameters. This approach ensures exhaustive coverage of all possible combinations of factor levels, facilitating a comprehensive analysis of their interactions and effects. However, the full factorial DoE generates all possible combinations of the levels of the independent factors, without considering any logical or technical constraints among configurations. In high-performance computing environments such as GROMACS, this can lead to execution failures, as certain combinations may be inconsistent or invalid with respect to the underlying hardware or the internal logic of the simulation software. For instance, combinations that assign more GPU tasks than physically available devices, or incompatible settings of the -nb, -pme, and -npme options, can result in runtime errors. A representative error message includes: *“There were N GPU tasks assigned on node A, but M GPU tasks were identified, and these must match.”* In practice, the full factorial design yielded both *successful* and *failed* configurations, depending on the compatibility of parameter combinations with the hardware and software constraints. The number of valid and invalid configurations varied across systems and benchmark cases. For the Villin system, 88 configurations completed successfully and 512 failed on Majorana, Franklin, and Ada, while 152 were successful and 2328 failed on Leonardo. For the RNase and ADH Dodecamer systems, the same trend was observed: 120 successful and 480 failed configurations on Majorana, Franklin, and Ada, and 200 successful versus 2,280 failed on Leonardo. Finally, for the STMV system, 120 configurations were successful and 480 failed on Majorana, Franklin, and Ada, while 232 completed successfully and 2248 failed on Leonardo. These results emphasize how, despite the exhaustive exploration provided by a full factorial DoE, only a small subset of the total combinations corresponds to valid configurations. The large proportion of failed runs reflects the presence of strong logical interdependencies among parameters in HPC environments such as GROMACS, where constraints related to GPU task allocation and internal domain decomposition often restrict the feasible configuration space.

To address this, combinations violating hardware constraints or software-specific logical requirements were identified and excluded from further analysis. This filtering step ensures that only feasible and meaningful configurations contribute to the performance evaluation, while preserving the methodological rigor of the factorial design.

### 2.5. Methodology: Automation of DoE and MD Simulations

All MD simulations were orchestrated through a custom workflow developed in Python 3.10. This environment allowed for seamless integration of experimental design, simulation execution, performance monitoring, and data post-processing. The workflow was structured to ensure full reproducibility and scalability across different GPU configurations.

To efficiently generate and manage the design space, we employed the *pyDOE3* library (v1.0.0), which facilitated the construction of a full factorial DoE matrix. Data handling and logging operations were managed using *pandas* (v1.5.3).

Notably, the settings for PME computation—namely *-npme*, *-pme*, and *-pmefft*—were fixed to values that consistently offloaded this calculation to the GPU. This choice was informed by preliminary screening experiments (data not shown), which demonstrated that GPU-based PME calculations outperformed CPU-based alternatives across all configurations tested. The finalized DoE matrix included 256 parameter combinations, which were exported as a CSV file (*gmx_fullfact_doe.csv*) to ensure transparency and traceability.

To guarantee consistency across runs, each simulation was initialized using unified topology (*.top*), structure (*.gro*), and parameter (*.mdp*) files, generating via gmx grompp the same MD configuration file (*.tpr*) for all the runs. The simulation runs were executed using *subprocess.run()* within Python, allowing fine-grained control over each call to *gmx mdrun*. This approach provided robust handling of standard output, standard error, and runtime exceptions, all of which were logged to a central file (*doe.log*) for auditing and debugging purposes.

The execution command for each simulation followed a common template:

*gmx mdrun -ntmpi X -pin Y -gputasks Z -bonded A -update B -nb gpu -pme gpu -pmefft gpu -npme 1 -nsteps 25000*

where the parameters *X*, *Y*, *Z*, *A*, and *B* varied according to the experimental design. The number of MD steps was fixed at 25000 to standardise throughput measurement, which was extracted from the GROMACS log file by parsing the line containing the “Performance” keyword. This throughput value (in ns/day) was appended in real time to the DoE matrix, allowing immediate performance comparison across configurations.

Upon completion of each simulation, temporary files were removed to minimize storage overhead, preserving only performance logs and GPU usage files (*gpu_usage_runX.csv*). The final results were consolidated into a structured dataset (*gmx_fullfact_doe_results.csv*) combining all experimental parameters with their corresponding throughput values.

Although the simulations did not rely on stochastic components, the use of identical input files ensured deterministic behaviour across executions. The total wall-clock time for each simulation was recorded and used as a reference for workflow scalability.

### 2.6. Data Analysis

Given that our response consists of continuous performance metrics, analysis of variance (ANOVA) is the preferred inferential tool for evaluating the significance of each factor and interaction term ^29,30^. ANOVA was employed to quantify the contribution of each experimental factor to the observed variability of the response variable and to assess whether the apparent effects were statistically distinguishable from background noise. The rationale behind this approach lies in the decomposition of the total variability of the system into systematic sources, attributable to controlled factors and their interactions, and random error, attributable to uncontrolled perturbations and measurement uncertainty. Formally, the model can be expressed as:

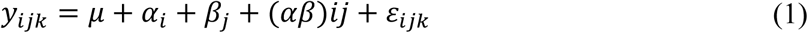

where *y*_*ijk*_ represents the observed response under the *i*-th level of factor *A*, the *j*-th level of factor B, and the *k*-th replicate. Here, μ denotes the overall mean, α_*i*_ and β_*j*_ correspond to the main effects of factors *A* and *B*, (αβ)_*ij*_ captures the interaction effect, and ε_*ijk*_ denotes the residual error assumed to be independently and normally distributed with constant variance. In the general DoE setting, the model can be extended to accommodate additional factors and higher-order interactions. The inferential procedure of ANOVA relies on partitioning the total sum of squares (*SSQ*_*tot*_) into additive components associated with each factor, their interactions, and the residual error. This decomposition is written as:

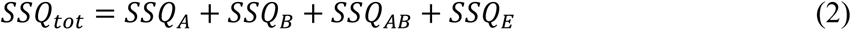

with each term subsequently normalized by its respective degrees of freedom to yield the mean square (*MS*). The central test statistic, the F-ratio, is computed for each factor or interaction as:

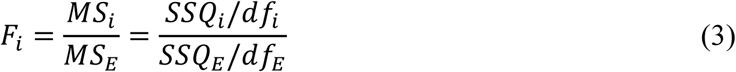

where *df*_*i*_ and *df*_*E*_ denote the degrees of freedom of the factor of interest and the error term, respectively. Under the *null hypothesis* that the factor has no effect, the distribution of *F*_*i*_ follows an F-distribution with (*df*_*i*_, *df*_*E*_) degrees of freedom, corresponding to the factor of interest and the residual error, respectively. A comparison between the observed F-statistic and the theoretical F-distribution enables the calculation of a p-value, which quantifies the probability of obtaining such an extreme ratio under the null hypothesis. Factors yielding p-values below the pre-specified significance threshold (typically α = 0.05) are considered statistically significant contributors to the response variability. More formally, given an observed value *F*^*obs*^_i_ for the *i*-th factor, the p-value is defined as:

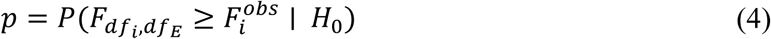

where *F*_*dfi*,*dfE*_ denotes a random variable following the F-distribution. This probability quantifies the degree of compatibility between the observed data and the null model of no effect.

The decision-making process in ANOVA is thus anchored in the comparison of the calculated p-value with a pre-established significance level α. If *p* < α, the null hypothesis is rejected, and the factor is deemed to exert a statistically significant influence on the response variable. Conversely, if *p* ≥ α, the null hypothesis cannot be rejected, and the data do not provide sufficient evidence to attribute systematic variation to the factor. It is important to underscore that the p-value does not quantify the probability that the null hypothesis is true; rather, it expresses the extremeness of the observed test statistic relative to the null distribution. In the specific setting of DoE, where multiple factors and interactions are simultaneously tested, the p-value serves as a critical criterion for discriminating between genuine effects and random fluctuations. When interpreted alongside effect sizes and confidence intervals, it ensures that the identification of significant parameters is both statistically rigorous and scientifically meaningful.

Within the DoE context, this ANOVA-based procedure is of critical importance for disentangling the relative importance of main effects and interaction terms. By systematically evaluating each source of variation, ANOVA provides not only a rigorous statistical test of factor relevance but also a foundation for model reduction and optimization strategies. This methodological framework ensures that the identification of significant parameters is not confounded by random fluctuations, thereby enhancing the robustness of subsequent predictive modelling. Overall, the integration of ANOVA into the DoE workflow enables the systematic exploration of the experimental space with statistical rigor. It transforms raw measurements into actionable insights, establishing whether variations in the studied response are the result of deliberate manipulations of input factors or merely stochastic noise. By doing so, it provides the methodological backbone for optimising factor settings and drawing robust scientific conclusions.

## 3. RESULTS AND DISCUSSION

The results of our study on the development and application of GROMODEX are presented below. For clarity of exposition, we present the results from a molecular-system-centric perspective. Readers are encouraged to refer to the Supplementary Information (SI), where the dataset is organized to vertically isolate each HPC system. This structure provides a clearer framework for analysing how each HPC system adapts to molecular systems of different sizes.

### 3.1 Performance Analysis

#### 3.1.1 Villin

Performance analysis of the simulations revealed substantial variability across runs (**Figure 1**). The dataset contains a significant proportion of executions yielding zero performance, indicating that these simulations did not execute successfully. The underlying causes can be attributed both to inconsistencies in the input files provided to GROMACS (see section **2.4**) and to the inability to apply a domain decomposition strategy involving a large number of MPI ranks on a system of limited size. In such cases, the overhead associated with managing excessive parallel tasks exceeds the computational benefits, resulting in aborted runs.

**Figure 1.**
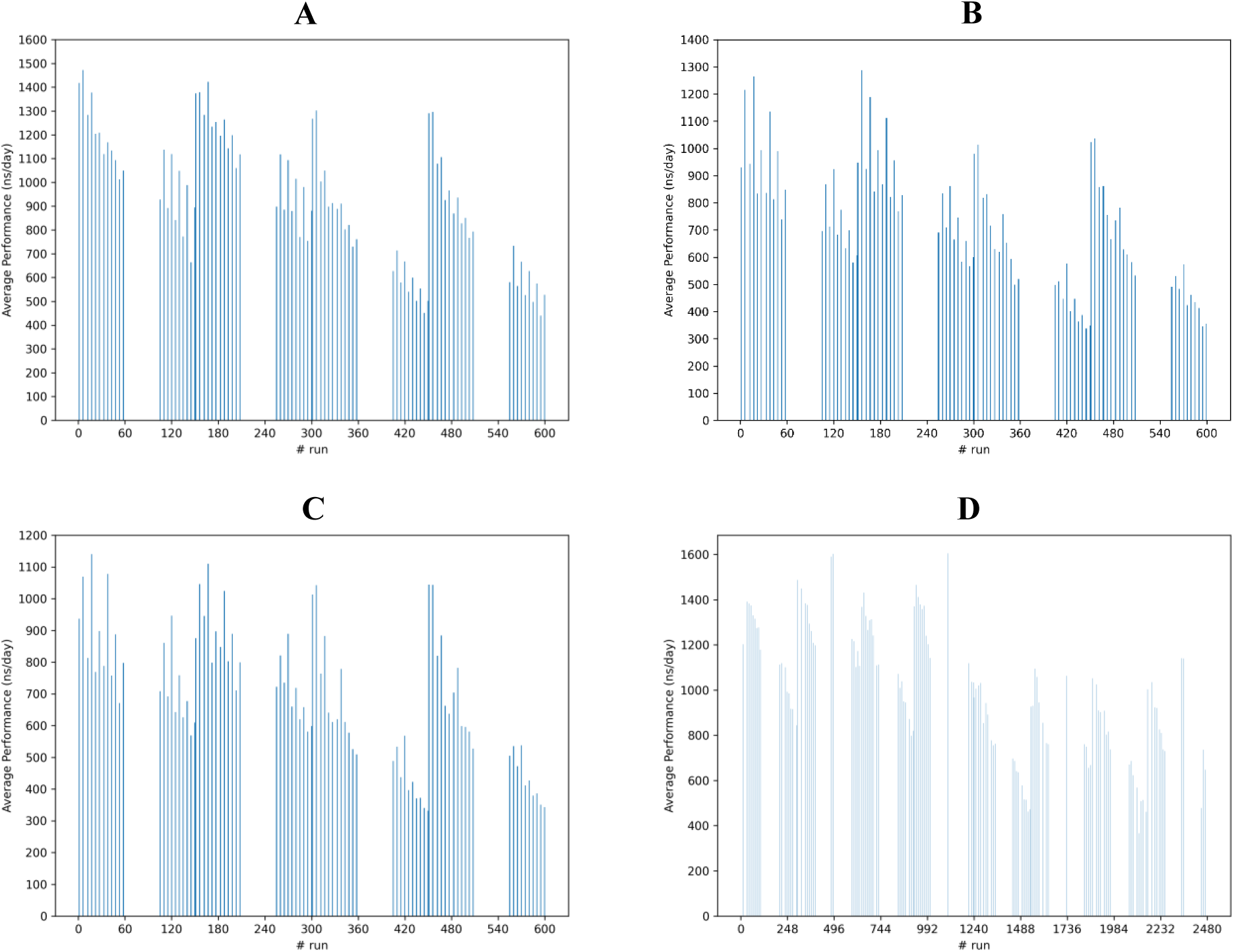
Average performance trend (three replicas for each HPC system) of Villin as a function of parameter settings. (A) Majorana, (B) Franklin, (C) Ada and (D) Leonardo.

**Figure 2.**
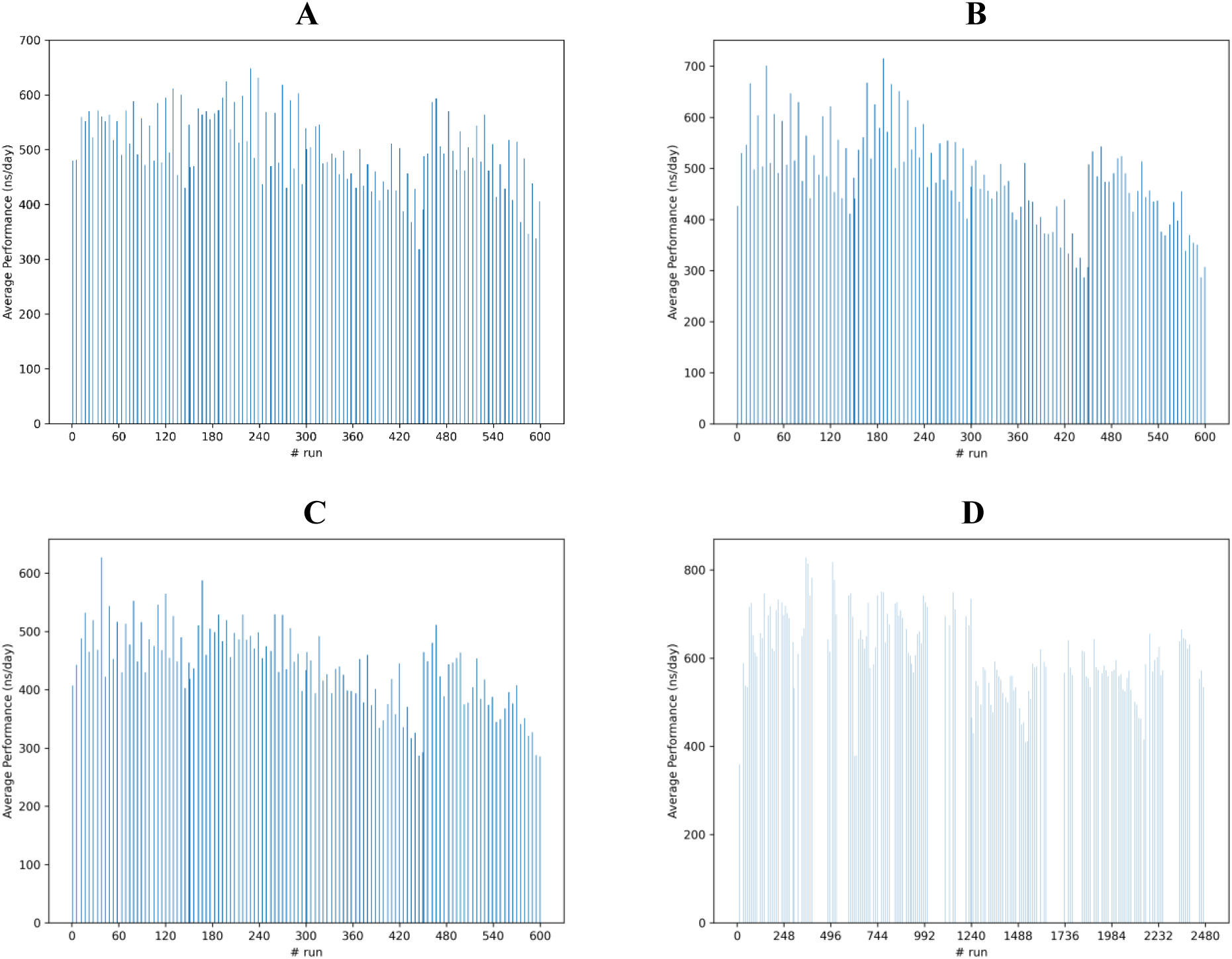
Average performance trend (three replicas for each HPC system) of RNAse Cubic as a function of parameter settings. (A) Majorana, (B) Franklin, (C) Ada and (D) Leonardo.

**Figure 3.**
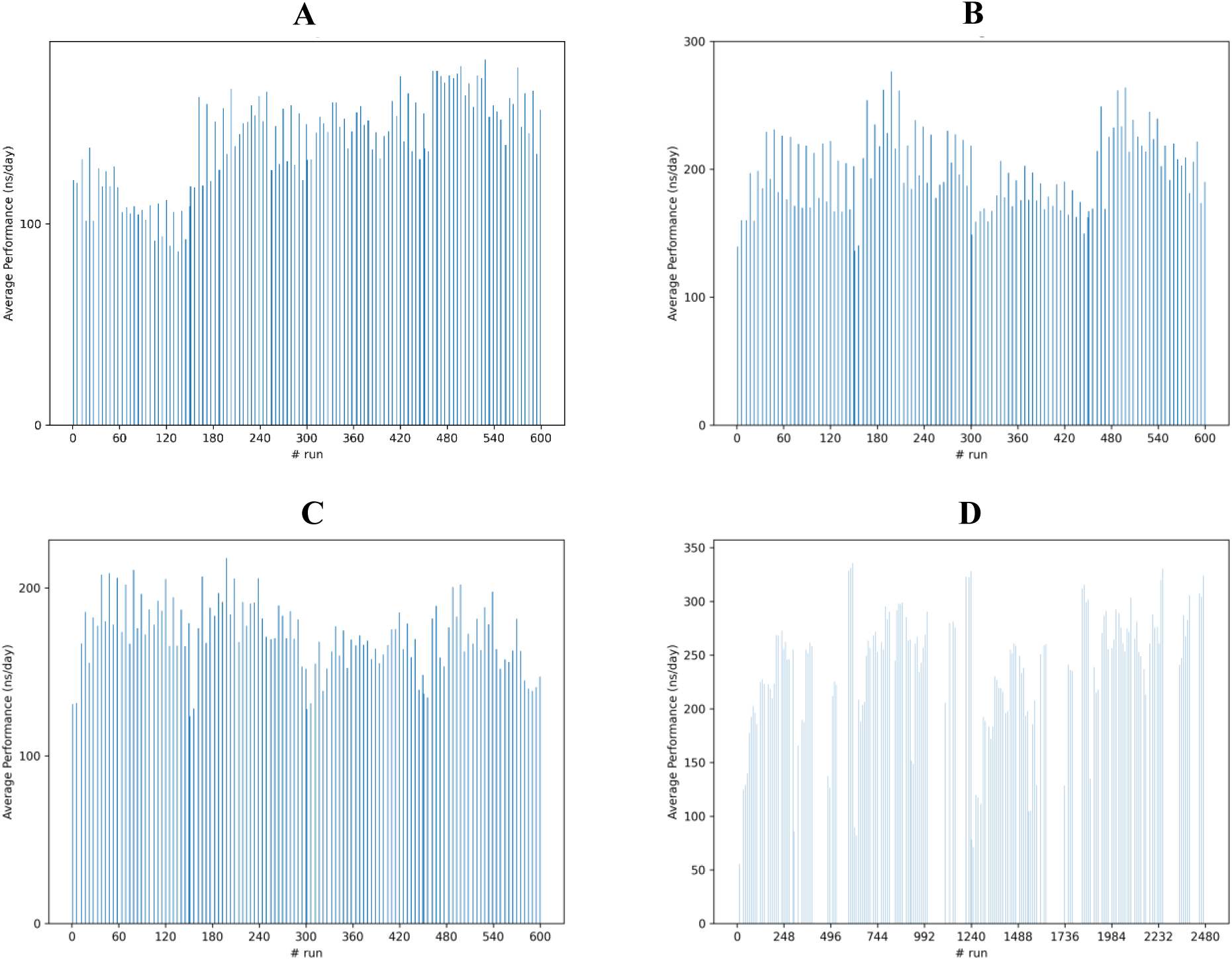
Average performance trend (three replicas for each HPC system) of ADH Dodec as a function of parameter settings. (A) Majorana, (B) Franklin, (C) Ada and (D) Leonardo.

Among the successful executions, performance data tend to cluster around specific index values, suggesting a discrete set of configurations with stable computational behaviour. Worth of note, the emergence of a sawtooth-like pattern in the overall trend; a preliminary visual inspection of this behaviour suggests that optimal performance is achieved at configurations with a relatively low number of MPI ranks, which allow for a more balanced load distribution and reduced communication overhead between processes.

**Table 3** summarises the parameters to which the highest performance corresponds for the Villin system across the four HPC configurations, revealing several non-trivial trends that cannot be directly ascribed to the underlying hardware characteristics. Leonardo achieves the highest simulation speed at 1605.220 ns/day, followed by Majorana (1471.406 ns/day), Franklin (1287.939 ns/day), and Ada (1139.921 ns/day). Despite its comparatively modest hardware configuration, Majorana outperforms both Franklin and Ada, securing second place in overall performance. This result can be partially explained by the hardware specifications: Leonardo is equipped with four custom NVIDIA A100 GPUs, offering not only higher memory bandwidth but also superior computational capabilities. In contrast, Majorana’s better-than-expected performance—despite having fewer and less powerful GPUs—may be attributed to its CPU’s high base frequency, which likely compensates through faster execution of CPU-bound tasks or more efficient GPU synchronisation. Leonardo’s CPU, while offering the highest thread and memory bandwidth potential, operates at a relatively moderate base clock (**Table 1**). Notably, Leonardo was the only system with *pin* set to off, suggesting that despite the lack of core binding, the GPUs resources compensate for any potential inefficiencies due to thread migrations. Conversely, the other systems-maintained *pin* on, likely to maximize CPU affinity and memory locality.

**Table 3.**
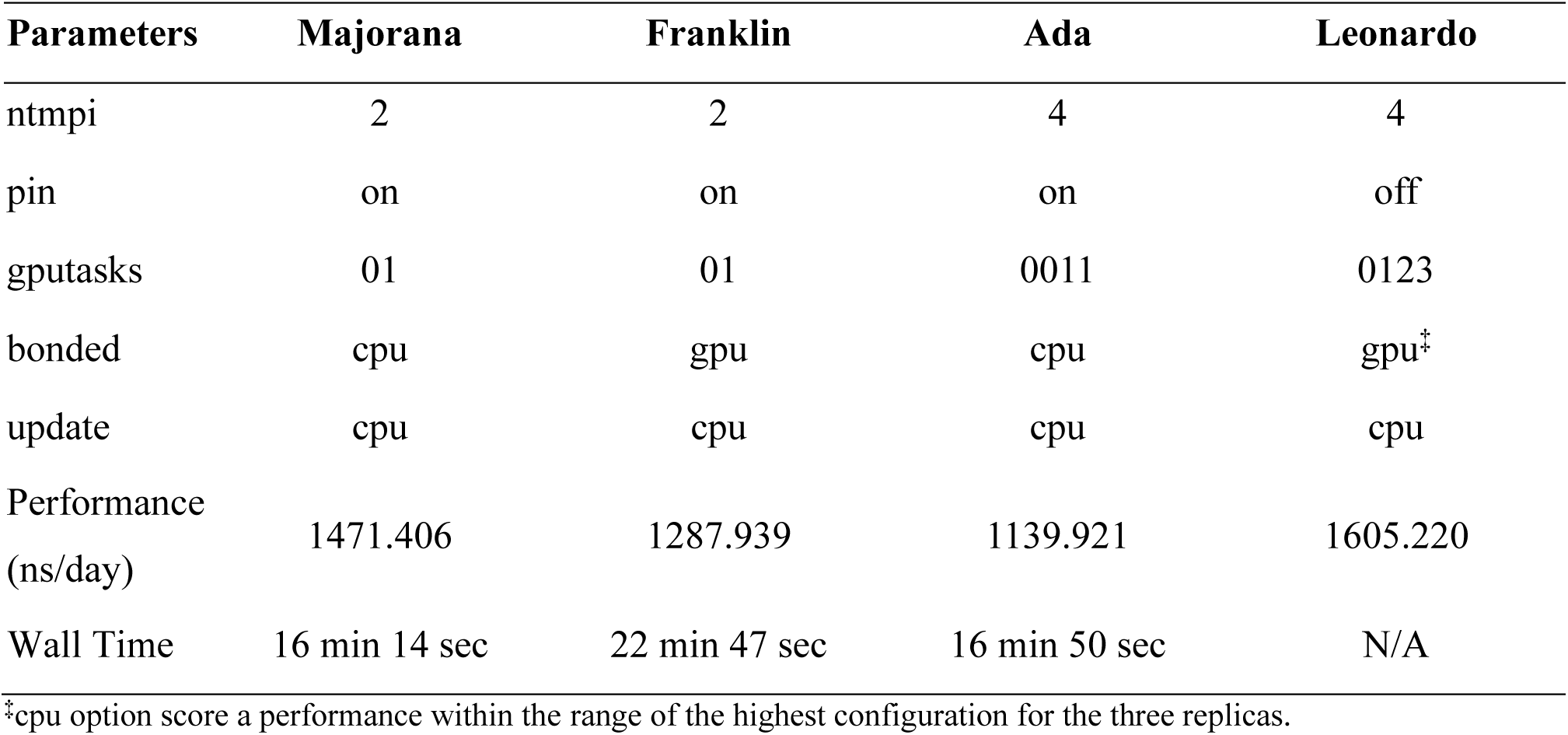
Highest performance parameter settings for Villin.

| Parameters | Majorana | Franklin | Ada | Leonardo |
| --- | --- | --- | --- | --- |
| ntmpi | 2 | 2 | 4 | 4 |
| pin | on | on | on | off |
| gputasks | 01 | 01 | 0011 | 0123 |
| bonded | cpu | gpu | cpu | gpu<sup>‡</sup> |
| update | cpu | cpu | cpu | cpu |
| Performance<br>(ns/day) | 1471.406 | 1287.939 | 1139.921 | 1605.220 |
| Wall Time | 16 min 14 sec | 22 min 47 sec | 16 min 50 sec | N/A |

<sup>‡</sup>cpu option score a performance within the range of the highest configuration for the three replicas.

The *gputasks* distribution further highlights architectural differences. On Majorana and Franklin both GPU were assigned to perform a single task each (i.e., 01). The same happened for Leonardo, which adopt a broader distribution of *gputasks* (i.e., 0123). In particular, the common feature of these two configurations lies in the fact that they both dedicate exclusively one of the available GPUs to the calculation of PMEs; in particular, in the GROMACS’ syntax, this is identified by the ID of the GPU present as the last value in the string (i.e., 1 for Majorana and Franklin, 3 for Leonardo). Conversely, for Ada the optimal performance was achieved with a 0011 *gputasks* configuration.

The allocation of *bonded* and *update* tasks is particularly interesting: while Majorana and Ada kept both tasks on the CPU, Franklin and Leonardo - which use the same type of GPU - instead offloaded the bonded calculations to the GPU. However, for Leonardo, a configuration with the bonded ones performed on the CPU still achieves near-peak performance. This strategy, especially in the case of Franklin, likely contributed significantly to reducing the computational load on the CPU, thus improving the overall throughput.

#### 3.1.2 RNAse Cubic

For the RNAse Cubic system, which is computationally more demanding, the performance results are noticeably more abundant in terms of successfully completed runs. This improvement is likely attributable to GROMACS’ ability to access to a higher number of MPI ranks in this context, thus expand the domain decomposition opportunities. The increased computational load appears to better justify the communication overhead, making parallelisation more efficient.

The highest performance values are observed in the mid-to-low region of the plots, although this peak emerges slightly later than in the case of the Villin system. In fact, as the dimensionality and complexity of the molecular system increase, HPC architectures tend to achieve optimal performance with different parameter settings, suggesting that alternative spatial decomposition strategies become more effective under these conditions. Moreover, the performance pattern for RNAse Cubic exhibits considerably greater uniformity, with fewer fluctuations and anomalies. Nonetheless, a consistent trend across the four HPC systems is the presence of a pronounced performance degradation at run index 445 for Majorana, Franklin, and Ada, and at index 1527 for Leonardo. This behaviour is associated with the configuration *ntmpi*: 10, *pin*: off, *gputasks*: 0000000001, *bonded*: cpu, and *update*: gpu, although this configuration does not necessarily correspond to the absolute minimum performance within the dataset.

As for the parameters with the best performance, Leonardo again leads with 828.215 ns/day, followed by Franklin (715.195 ns/day), Majorana (648.367 ns/day), and Ada (627.306 ns/day), as can be seen from **Table 4**. The relative performance advantage of Majorana is less pronounced here with respect to the Villin case, possibly indicating a scaling limitation when system size increases and communication overhead becomes more relevant.

**Table 4.** Highest performance parameter settings for RNAse Cubic.

| Parameters | Majorana | Franklin | Ada | Leonardo |
| --- | --- | --- | --- | --- |
| *ntmpi* | 8 | 6 | 6 | 9 |
| *pin* | on | on | on | off* |
| *gputasks* | 00000111 | 000111 | 000111 | 000111222* |
| *bonded* | gpu | gpu | cpu | cpu |
| *update* | cpu | cpu | cpu | cpu |
| Performance<br>(ns/day) | 648.367 | 715.195 | 627.306 | 828.215 |
| Wall Time | 28 min 10 sec | 34 min 36 sec | 29 min 15 sec | N/A |

Regarding *ntmpi*, Majorana required more MPI ranks than Ada and Franklin, likely due to its weaker hardware.

All systems except Leonardo used *pin* set *on*. The fact that Leonardo again achieved the best performance with *pin* set *off* suggests that in this particular configuration binding cores can sometimes become a bottleneck rather than a benefit, especially when GPU communication and synchronization dominate runtime.

The *gputasks* mappings further explain the performance patterns: Leonardo distributed *gputasks* across three GPUs (000111222). Majorana, Franklin, and Ada limited the load to two GPUs (000111), with respect to Leonardo. Interestingly, Majorana adopted a non-symmetrical distribution of *gputasks*, preferring to allocate more computing resources to PME on GPU #1.

In all cases bonded and update tasks were computed on the CPU, suggesting a consistent design strategy to balance the CPU–GPU workload.

#### 3.1.3 ADH Dodec

Performance analysis for the ADH Dodec system, shown in Figure 3 Table ***5***, exhibits a complex and heterogeneous behaviour, reflecting the computational demands of this larger biomolecular system. In contrast to the more compact systems previously analysed, ADH Dodec displays a broader dynamic range in performance values, as well as wider sensitivity to the configuration parameters adopted in each run.

The benchmarking results for the ADH Dodec system, a significantly larger molecular system, illustrate once again the critical role of hardware architecture and task distribution in determining GROMACS performance (**Table 5**).

**Table 5.** Highest performance parameter settings for ADH Dodec.

| Parameters | Majorana | Franklin | Ada | Leonardo |
| --- | --- | --- | --- | --- |
| *ntmpi* | 8 | 6 | 6 | 16 |
| *pin* | on | on | on | on |
| *gputasks* | 00000111 | 000011 | 000011 | 0000011111222223 |
| *bonded* | gpu | gpu | gpu | cpu |
| *update* | gpu | cpu | cpu | cpu |
| Performance<br>(ns/day) | 181.478 | 276.369 | 217.668 | 340.213 |
| Elapsed Time | 1 h 15 min 25 sec | 1 h 3 min 48 sec | 1 h 3 min 7 sec | N/A |

Leonardo outperforms all other platforms with a simulation speed of 340.213 ns/day, followed by Franklin (276.369 ns/day) and Ada (217.668 ns/day), while Majorana suffered of poor computational capabilities (181.478 ns/day), almost losing half the performance compared to Leonardo.

Regarding parallelization, Leonardo utilized 16 MPI ranks, almost three time the number used on Franklin and Ada (6 each) and twice that of Majorana (8). The higher *ntmpi* setting on Leonardo facilitated better domain decomposition and load balancing across the multiple GPUs and CPU cores, essential for efficiently handling such a large molecular system.

The *pin* parameter was uniformly set to *on* across all platforms for this benchmark, reflecting the necessity of strict CPU-GPU affinity to minimize memory access latency and optimise interconnect usage under heavier computational loads.

A deeper look into the *gputasks* assignment showed substantial common features. At this level of dimensionality of molecular systems, a non-symmetrical allocation of GPU tasks was most effective. In fact, Majorana, Franklin and Ada have adopted configurations for this parameter that leave a large part of GPU #1 dedicated to the calculation of PMEs, which, at this level of dimensionality, constitute an important part of the computational load. Leonardo, on the strength of its configuration, distributed the workload across all four GPUs with finer granularity, dedicating one GPU (ID: 3) to exclusively PME computation.

The assignment of *bonded* and *update* tasks also presented an interesting deviation. While Majorana, Franklin, and Ada offloaded *bonded* interactions to the GPU to reduce CPU bottlenecks, Leonardo processed *bonded* interactions on the CPU, without penalizing GPU workload.

#### 3.1.4 STMV

The performance evaluation of the STMV system highlights a progressive and well-structured pattern. As a considerably large and complex biomolecular system, STMV presents consistent trends that reflect how hardware is challenged under such intensive demand of computational resources.

Figure 4A displays a well-defined staircase-like increase in performance, with three clearly distinguishable regions of growth, suggesting strong saturation in computational scaling. These plateaus, less identifiable in other molecular systems under consideration, fall at a specific combination of the *bonded* and *update* levels that unlock higher throughput. The steep performance increase around runs 450 indicates a jump in efficiency, due to progressive computational offload of the GPUs at those configurations.

**Figure 4.**
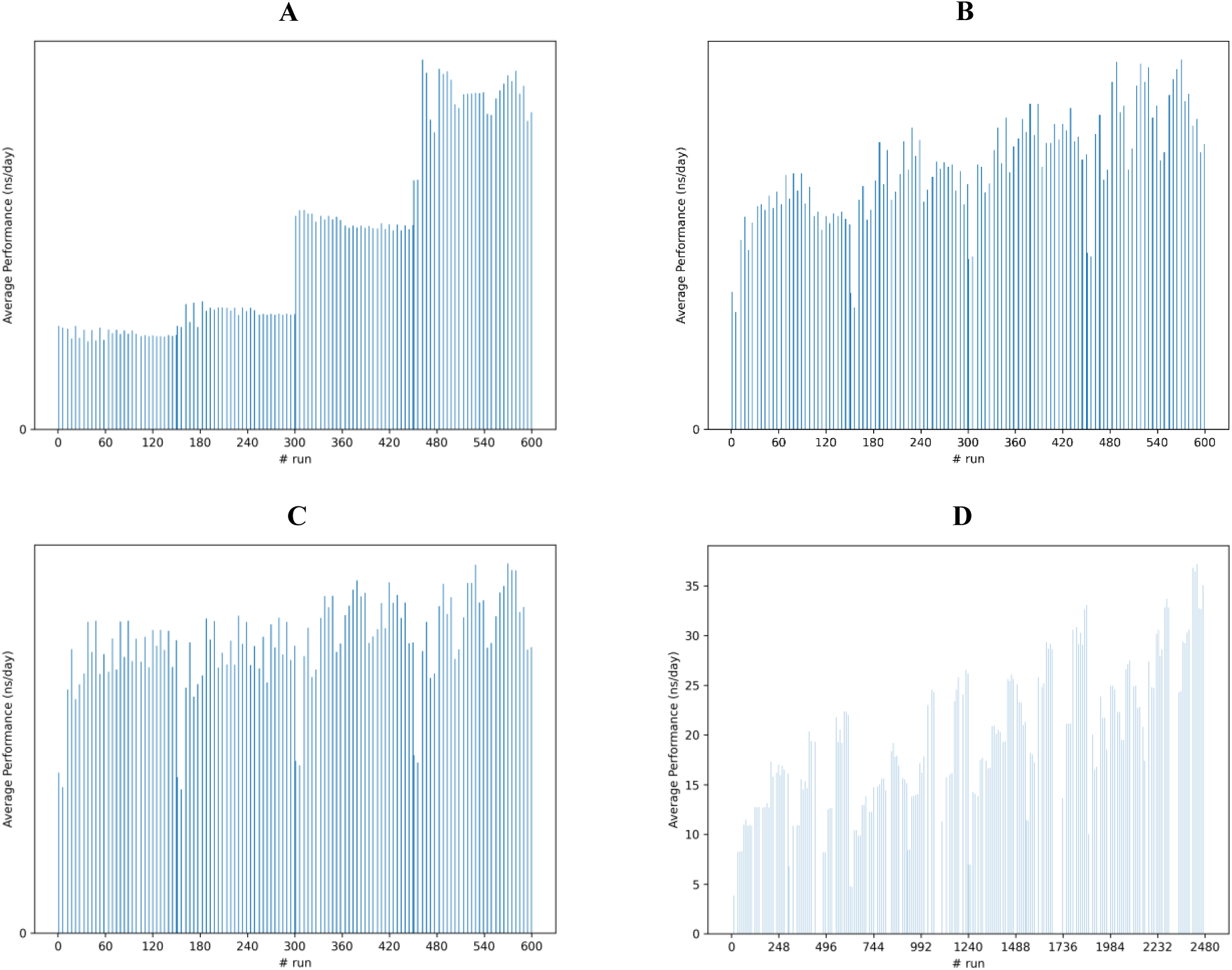
Average performance trend (three replicas for each HPC system) of STMV as a function of parameter settings. (A) Majorana, (B) Franklin, (C) Ada and (D) Leonardo.

Both Franklin and Ada exhibit a trend of gradual performance improvement characterized by multiple local peaks (Figure 4B and **4C**). This indicates a nuanced scaling behaviour in which many configurations yield similar, though subtly distinct, performance scores. In Franklin’s case, the lack of sharp drops may point to better load balancing or greater system stability, while Ada shows more noticeable variability across individual runs. This variability suggests that, despite the overall improvement, performance remains sensitive to specific configuration choices—highlighting the importance of fine-tuning parallel parameters even within comparable scaling regions.

Leonardo in Figure 4D, covering a broader range of runs, reveals a consistent upward trend, confirming a strong sensitivity in performance over a wide range of configurations. This case study corroborates more than ever that, despite mere computing power, the sub-optimal choice of simulation parameters can lead to a drastic degradation of simulation performance, especially in contexts where the molecular system is particularly challenging.

Leonardo once again demonstrated superior performance with a simulation speed of 37.197 ns/day, followed by Franklin (26.687 ns/day), Ada (19.745 ns/day), and, considerably behind, Majorana (10.740 ns/day). The results are once again summarised in **Table 6**.

**Table 6.**
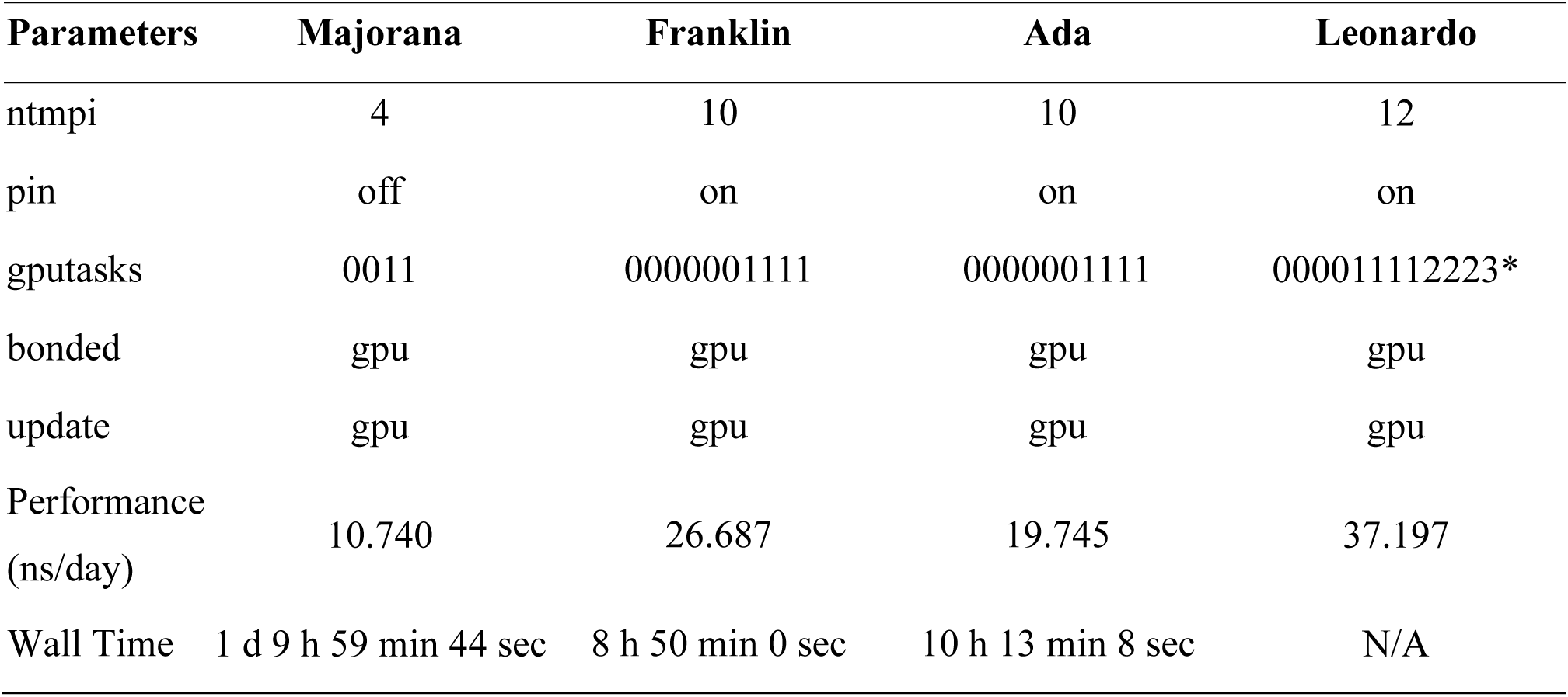
Highest performance parameter settings for STMV.

Notably, the performance gap widens significantly at this scale, particularly between Majorana and the other systems. Majorana’s relatively modest hardware proved inadequate to sustain competitive throughput. The lower *ntmpi* on Majorana clearly constrained its ability to decompose the large system into enough parallel tasks, severely bottlenecking performance. By contrast, the higher *ntmpi* on Franklin, Ada, and Leonardo enabled a much finer domain decomposition, essential for simulations with such a massive number of atoms.

The configuration of CPU affinity represents a key differentiating factor in system performance across platforms. On systems such as Franklin, Ada, and Leonardo, CPU affinity was explicitly enabled (*pin on*), a choice that facilitated optimised memory access patterns and reduced cache contention, thereby contributing to overall computational efficiency. In contrast, Majorana operated without CPU pinning (*pin off*), a setting that further magnified its performance limitations under intensive computational workloads.

In terms of *gputasks* assignment, all systems employed non-symmetrical distribution strategies akin to those observed in ADH Dodec. Franklin and Ada deployed a mapping scheme spanning ten GPU tasks (0000001111), reflecting a balanced distribution over two physical GPUs. Leonardo advanced this approach with a more granular and extended mapping scheme (000011112223), with one GPU dedicated to PME calculation only. In contrast, Majorana, constrained by a limited computational capacity, relied on a simpler allocation pattern (0011), lacking the granularity and scalability required for efficiently support the simulation effort.

Interestingly, for STMV, all systems offloaded both *bonded* and *update* calculations to the GPU, a reflection of the critical need to minimize CPU overhead at such a scale; even Leonardo kept bonded interactions on the GPU here, recognizing that CPU processing would be inefficient compared to the vast parallel capabilities of its GPUs for such a large problem size.

#### 3.1.5 Right-sizing

Before proceeding with the statistical analysis, it is useful to consolidate the performance results to highlight the underlying trends across architectures and workloads. This section serves to reframe the data from a broader perspective, emphasizing how computational efficiency emerges from the balance between system size and hardware characteristics. Figure 5 summarises the performance obtained across the four HPC architectures for the four molecular systems. Each curve represents one system, showing how performance scales with the architectural configuration.

**Figure 5.**
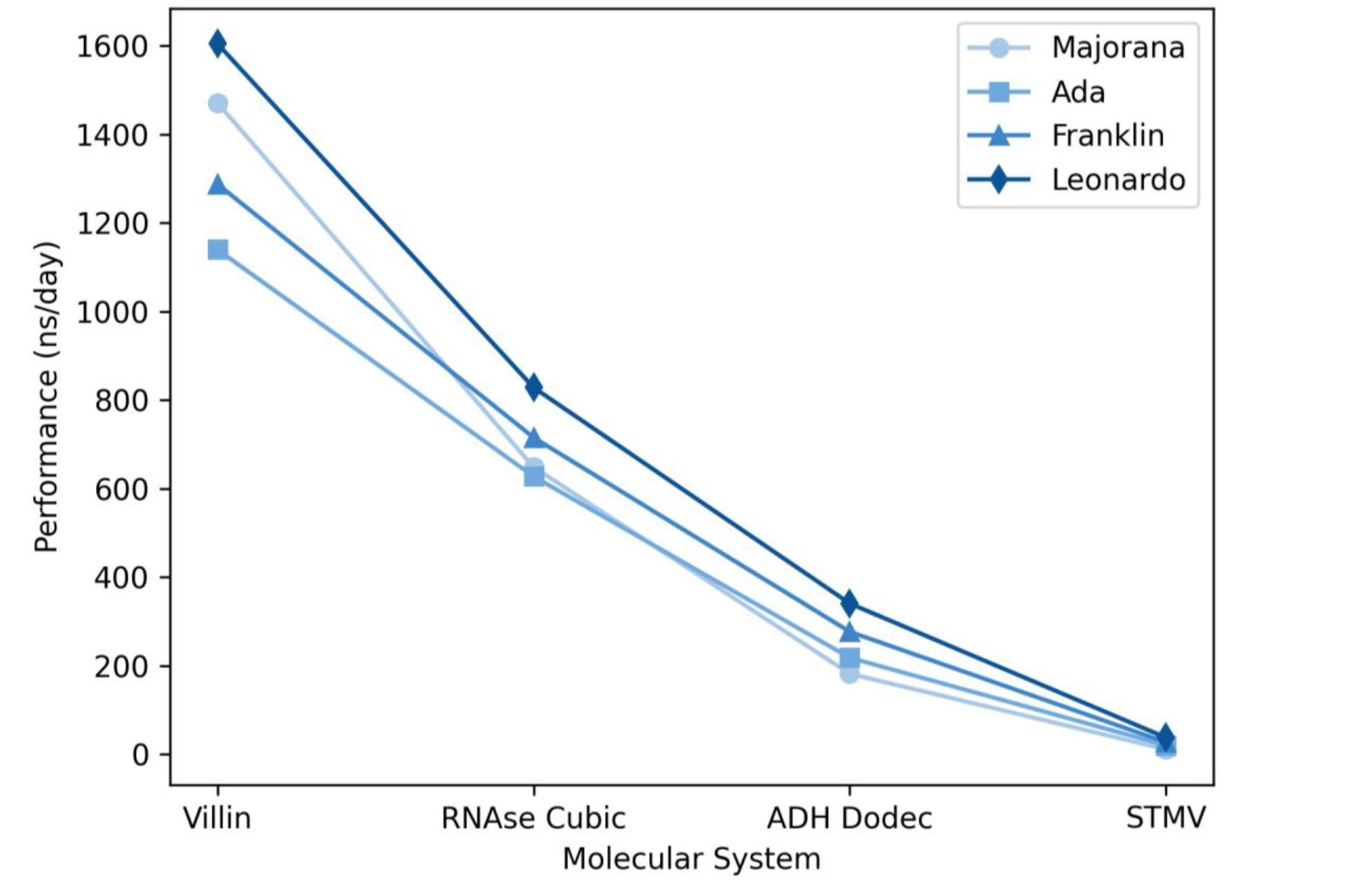
Performance comparison across four HPC architectures for the four molecular systems considered. Each curve represents a distinct system, illustrating how computational efficiency scales with architectural characteristics. The figure highlights the right-sizing effect, where optimal performance emerges from the alignment between workload scale and hardware resources.

Majorana outperforms larger HPC systems for small-scale molecular dynamics, particularly in the Villin case. This behaviour can be attributed to the limited degree of parallelism involved, which allows the i9 CPU to exploit its higher clock frequency (**Table 1**) and achieve superior per-thread efficiency. In contrast, the other systems (RNAse, ADH, STMV) exhibit improved scalability on architectures with broader parallel resources, where communication overhead becomes less dominant relative to the total computational workload. These findings indicate that no universally optimal configuration exists; rather, performance depends on a dynamic interplay between workload characteristics, system size, and architectural design. This *right-sizing* perspective provides a conceptual bridge to the following ANOVA, which quantifies the statistical significance of the factors influencing performance.

### 3.2 Main Effects and Interactions

Two-way ANOVA results (Figure 6Figure 6) across the four high-performance computing (HPC) architectures demonstrate a consistent pattern of both dominant main effects and critical interaction terms influencing computational performance across workloads. Despite architectural and workload-specific variability, the *gputasks* and the *ntmpi* systematically emerge as the most impactful parameters shaping execution efficiency. On all platforms, these two factors exhibit exceptionally low p-values (p ≪ 0.001), achieving statistical significance and accounting for a substantial portion of the variance in all simulation performance. This consistency highlights the universal importance of GPU workload partitioning, confirming that GPU-level parallelism remains a core determinant of performance across both cutting-edge and legacy architectures. *ntmpi* often surpasses *gputasks* in significance, particularly on architectures such as Leonardo and Franklin, where MPI process distribution critically shapes data locality and interprocess communication overhead. In particular, *ntmpi* achieves particular relevance in those situations where exceptional sensitivity of these architectures to the granularity of parallel decomposition (**Table 5** and **Table 6**). These results not only reaffirm the central role of hardware-aware parallel configuration – i.e., configurations that explicitly consider the system’s architecture such as core counts, memory hierarchy, and CPU-GPU topology – but also stress the need to tailor process-level and thread-level resource allocation strategies to system-specific architectural constraints.

**Figure 6.**
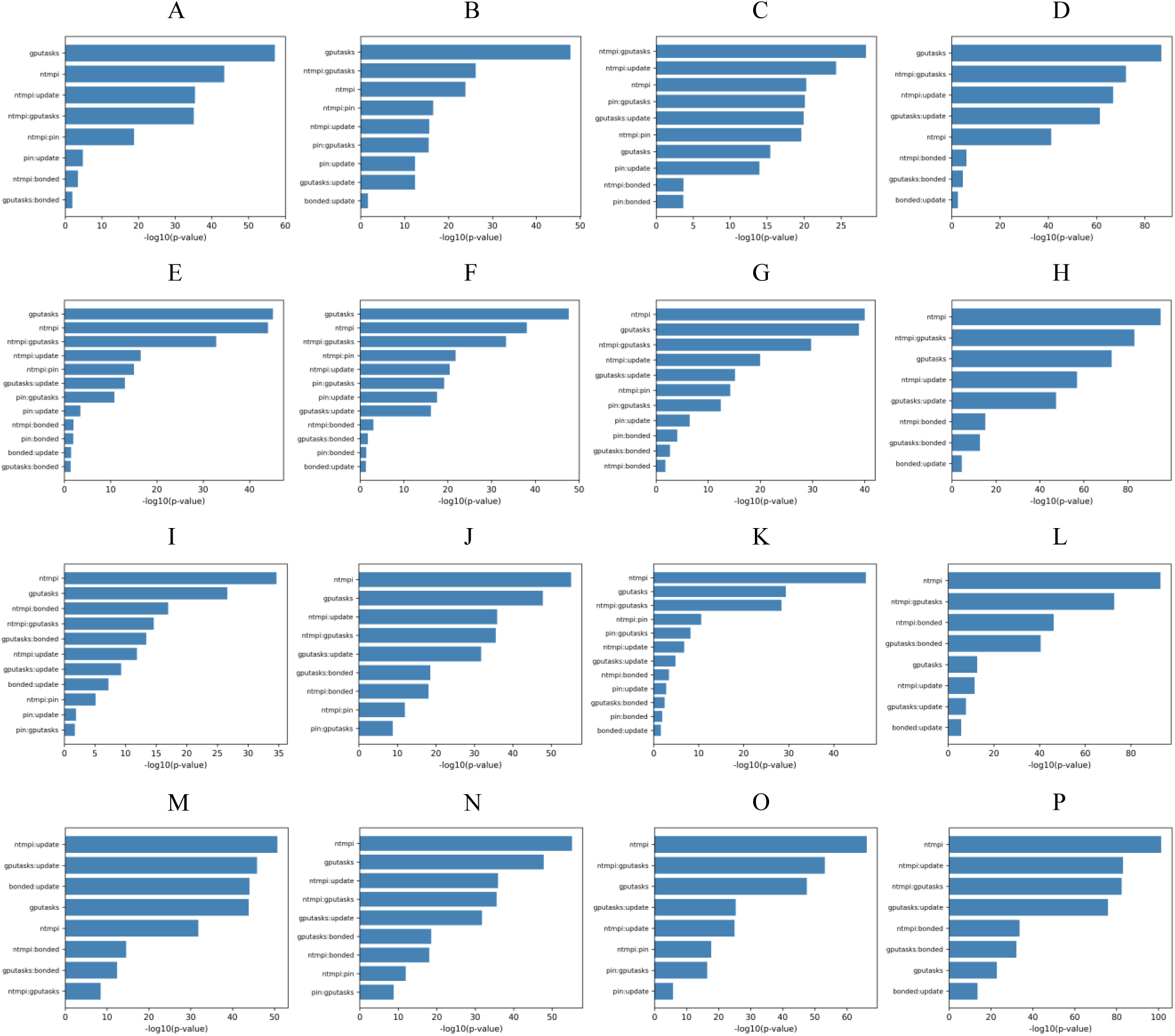
Two-way ANOVA results: the higher the bar, the more relevant the factor – or combination of two factors – is. From top: Villin, RNAse Cubic, ADH Dodec, STMV. From left: Majorana, Franklin, Ada, Leonardo.

Beyond individual factor effects, interaction terms provide key insight into the complex, non-linear nature of performance scaling in heterogeneous systems. Among these, the *ntmpi:gputasks* interaction repeatedly demonstrates significance across all systems, reinforcing that computational performance cannot be optimised by tuning MPI or GPU parameters in isolation. Instead, a synergistic multidimensional (or multifactorial) adjustment is necessary; this interaction captures a critical performance contour that reflects the load balancing and task scheduling challenges intrinsic to modern HPC environments.

Moreover, interactions involving variables that are excluded from the final models as individual factors—specifically *pin*, *update*, and *bonded*—consistently retain statistical significance in combination with *ntmpi* or *gputasks*. While their main effects yield p-values well above the 0.05 threshold, rendering them individually non-significant, these variables nonetheless participate in key second-order interactions that modulate performance outcomes. For example, the *pin:update* interaction displays significant effects on multiple circumstances, suggesting that memory affinity settings influence performance predominantly through their effect on *update* tasks or data locality during time integration steps. Similarly, *bonded:update* and *gputasks:bonded* achieve significance, illustrating that short-range interactions are sensitive to resource allocation primarily in the context of other configuration parameters. These observations underscore an important methodological insight in performance modelling: statistical insignificance of a factor in isolation does not equate to practical irrelevance. Instead, the exclusion of such factors from the main effects may suggest evidence of conditional importance^31^—a phenomenon in which the influence of a parameter becomes apparent only when coupled with others. This is especially relevant in high-dimensional configuration spaces, such as HPC performance tuning, where emergent properties often result from the interplay between multiple control variables. The relevance of these interaction terms varies not only across HPC platforms but also across workloads. For instance, *pin*, despite lacking individual significance, exhibits crucial interactions, like *ntmpi:pin* and *pin:bonded*, which affect memory access and CPU-GPU synchronisation, confirming the conditional relevance. Additionally, *bonded* and *update*, although often contributing less to overall variance when considered individually, become more relevant when considered in combination with *ntmpi* or *gputasks*. This finding is evident in statistically significant interactions such as *ntmpi:bonded* and *gputasks:bonded* on Leonardo, which indicate that the performance sensitivity of bonded calculations is largely determined by their GPU distribution. Across all systems, the explanatory power of the models is uniformly high, with residual variances remaining consistently low. This indicates that the selected main and interaction effects capture the majority of performance determinants and that the factorial design provides a robust framework for exploring configuration sensitivity. However, the findings also reveal that no single configuration strategy is universally optimal across architectures. Instead, the relative importance of factors and interactions shifts depending on HPC-molecular system pair characteristics.

## 4. CONCLUSIONS

This study presented the development and application of GROMODEX, a robust tool for systematically optimising GROMACS MD simulations through a DoE approach. By combining rigorous full-factorial experimental design, high-throughput automation and ANOVA, GROMODEX facilitates the exploration of complex parameter spaces to identify configurations that maximize simulation performance. The results underscore the versatility of the tool and its potential for advancing MD research.

It is important to note that the configurations explored in this study were primarily assessed in terms of computational performance and successful execution, rather than long-term physical stability or numerical reproducibility of the molecular dynamics trajectories. Certain parameter combinations—especially those affecting domain decomposition, thread affinity, and GPU task allocation—may influence not only performance but also simulation stability and numerical accuracy. Therefore, the results presented here should be interpreted as indicative of computational efficiency within the tested setups, while additional verification would be required to ensure full physical consistency of the simulations across different configurations.

The investigation of four molecular systems across four computational architectures (Majorana, Franklin, Ada and Leonardo) revealed critical insights into the interplay between system– specific requirements and hardware capabilities. The diversity of molecular systems ensured the generalizability of the adopted protocol, while the variation in computational platforms provided a nuanced understanding of architecture-specific optimisation strategies.

A key outcome of this work was the identification of optimal parameter configurations for each molecular system and architecture. Factors such as *ntmpi* and *gputasks* were shown to have significant impacts on simulation throughput, as evidenced by the two-way ANOVA. These findings highlight the necessity of tailored optimisation approaches that account for both molecular complexity and hardware specifications.

Beyond the identification of optimal configurations, the study provided a deeper understanding of the parameter interdependencies that influence performance. The interaction effects revealed by the ANOVA collectively demonstrate that performance optimisation on modern HPC systems cannot rely solely on analysis of primary factors. Instead, attention must be paid to the conditional and often non-additive nature of parameter interactions. While *ntmpi* and *gputasks* universally dominate, their impact is magnified or modulated by interactions with seemingly secondary variables such as *pin*, *update*, and *bonded*. Therefore, a comprehensive, interaction-aware approach is essential to unlock the full potential of computational resources. Neglecting such interactions risks oversimplifying the optimisation process and may result in configurations that are suboptimal or misleadingly efficient only under restricted conditions. These findings advocate for the integration of multifactorial analysis methods into performance tuning workflows, promoting a deeper understanding of system behaviour and enabling more informed decisions in production-scale simulations.

The adoption of tools like GROMODEX systematic and scalable approach addresses a critical need in the field, enabling researchers to achieve superior performance across diverse systems and architectures. The insights gained from this study not only validate the efficacy of GROMODEX but also provide a foundation for future developments in simulation optimisation, paving the way for more efficient and accurate MD research.

## ACKNOWLEDGMENTS

This research was supported by the Italian Research Center on High Performance Computing, Big Data and Quantum Computing (CN1-HPC) and the Italian SuperComputing Resource Allocation (ISCRA code: HP10C31W4Y).

## 5. DATA AVAILABILITY STATEMENT

The GROMODEX source code is available at: https://github.com/MarcoSavioli/GROMODEX.git.

